# RNA folding using quantum computers

**DOI:** 10.1101/2021.05.27.446060

**Authors:** Dillion M. Fox, Christopher M. MacDermaid, Andrea M.A. Schreij, Magdalena Zwierzyna, Ross C. Walker

## Abstract

The 3-dimensional fold of an RNA molecule is largely determined by patterns of intramolecular hydrogen bonds between bases. Predicting the hydrogen bonding network from the sequence, also referred to as RNA secondary structure prediction or RNA folding, is a nondeterministic polynomial-time (NP)-complete computational problem. The structure of the molecule is strongly predictive of its functions and biochemical properties, and therefore the ability to accurately predict the structure is a crucial tool for biochemists. Many methods have been proposed to efficiently sample possible secondary structure patterns. Classic approaches employ dynamic programming, and recent studies have explored approaches inspired by evolutionary algorithms. This work demonstrates leveraging quantum computing hardware to predict the secondary structure of RNA. A Hamiltonian written in the form of a Binary Quadratic Model (BQM) is derived to drive the system toward maximizing the number of base pairs while simultaneously maximizing the average length of the stems. An Adiabatic Quantum Computer (AQC) is compared to a Replica Exchange Monte Carlo (REMC) algorithm programmed with the same objective function, with the AQC being shown to be highly competitive at rapidly identifying low energy solutions. The method proposed in this study was compared to three algorithms from literature and was found to have the highest success rate.

## Introduction

RNA molecules fold into complex secondary structures, which determine many aspects of RNA function as well as molecular properties such as thermal stability and compactness. Thus, RNA folding has an impact on protein translation, transcriptional regulation, and other processes vital to cellular functions (1–3). Secondary structure is also key to the function of synthetic RNAs which are used in a variety of applications ranging from protein design and genome editing to mRNA vaccine development (for example, (4–6)).

Methods for RNA structure determination are therefore of great interest and importance for basic research, applied biotechnology, and rational drug discovery. Experimental approaches developed for this purpose are extremely time consuming and expensive, and therefore their use is limited in practice. Computational methods are an attractive alternative as they aim to predict the folded structure of an RNA molecule based solely on sequence information, which can be readily obtained from high-throughput sequencing experiments. There are varied approaches to computational RNA structure prediction, ranging from physics-based methods that assign thermodynamic scores to a pre-defined set of structural features (7–9), to deep learning models trained on large RNA databases (10–13).

Similar to proteins, secondary structure of RNA molecules is largely determined by the sequence. Unlike proteins, where the folding is a global process largely driven by hydrophobic forces, single-stranded RNA molecules undergo a hierarchical folding process that is dominated by the formation of hydrogen bonds between nucleotides. Compared to protein folding, the formation of RNA tertiary structure is relatively slow. The RNA folding process involves the formation of strong restrictive local geometries which usually results in well-defined, thermally stable structures with little flexibility compared to protein structures (14).

The standard RNA hydrogen bonding pairs, (GC and AU) are known as the Watson-Crick base-pairs. Another common type of interaction that can form, known as the Wobble interaction, is between G and U. A variety of local structures, such as internal loops, hairpin loops, stacks, bulged loops and multi-loops, can be formed by Hoogsteen- or CH-edges and Sugar-edges. Such edges have the ability to accommodate types of interactions other than Watson-Crick type edges, including the formation of base triplets (base-pairs between three bases) that can modulate the stability of helices all the way to quaternary structures. A set of consecutive base pairs is often referred to as a *stem*. RNA structures also undergo formation of long-range interactions such as pseudoknots, which occur when base pairs cross without overlapping.

RNA structure prediction is a computationally expensive task, particularly when the solution space includes pseudoknots. Folding algorithms that do not account for pseudoknots tend to scale polynomially, with the most efficient method having sub-cubic scaling (15). There also exist approximate methods that have been shown to have linear scaling (16). A widely used minimum free energy (MFE) approach to RNA secondary structure with pseudoknots is an NP-complete computational problem (17). For an RNA strand with *N* stems, there are a total of *2*^*N*^ possible combinations of stems that could define the folded structure. For example, the SARS-CoV2 spike glycoprotein segment of the Pfizer-BioNTech COVID-19 vaccine (18) contains approximately 486,000 possible stems, so the total combinatorial space contains 2^486,000^ (~ 10^146,440^) possible solutions. For reference, the number of atoms in the universe is approximately 10^80^, inferred from the cosmological parameters presented in the Planck Collaboration (19). Although many of the possible combinatorial solutions likely contain overlapping stems, the majority of the potential stems cannot be excluded from the set *a priori*. Therefore, the total number of evaluations required to exhaustively sample the solution space grows exponentially with the number of stems. Since the solution space cannot be exhaustively sampled for many biologically relevant RNA sequences, either an alternative approach is required to identify an approximate solution, or pseudoknots must be disregarded. Classic approaches to sampling RNA secondary structure configurations utilize dynamic programming (9,20,21), while more recent methods have employed simulated annealing and Monte Carlo methods (22–24). In this study, we investigate the viability of utilizing quantum computers (QCs) to efficiently identify high-quality solutions to RNA secondary structure prediction.

To date state-of-the-art quantum devices are able to outperform classical computers in a narrow range of tasks (25,26), however, there have not yet been any concrete demonstrations of quantum advantage for commercial applications primarily due to the fact that QCs are limited in size and capabilities (27). It is speculated that the pharmaceutical, chemical, and life sciences industries will be the first fields to benefit from quantum computing technology (28,29). While applications utilizing quantum mechanical calculations, such as quantum chemistry, have clear maps to QCs (see, for example, (30–33)) at present these problems require too many qubits for current hardware to solve industrially relevant problems (29,34). To date there are primarily two models of QCs: the gate-based and the Adiabatic Quantum Computer (AQC). Gate-based quantum devices have a broad application range and are the most commonly used for quantum chemistry and quantum machine learning calculations. An alternative design, pioneered by the company D-Wave, is the AQC. Compared to the multitude of applications of gate-based QCs, AQCs have a much narrower range and only focus on optimizing solutions to problems by minimizing a problem Hamiltonian. To date, AQCs containing thousands of qubits have been built, and these devices are capable of solving sufficiently large discrete combinatorial optimization problems to permit testing against real world industrial use-cases. Rather than being programmed by sequences of quantum operators, AQCs are designed to anneal quadratic Hamiltonians in the form of Equation (1).

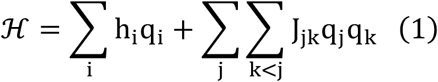

Here, *q*_*i*_, *q*_*j*_, and *q*_*k*_ represent the values of the qubits which can either be {0, 1} or {−1, 1} for binary or spin representations, respectively, *h*_*i*_ are the one-body terms, and *J*_*jk*_ are the two-body interactions. For the Ising model of a ferromagnet, *h* represents the magnetic dipole moments of the atomic spins and *J* represents the energy of the interactions between the spins. The D-Wave AQC used in this study is an analog device containing approximately 5,000 qubits. The device is programmed by setting values of local magnetic fields and coupling strengths, and the annealing process works by adiabatically lowering the strength of a transverse magnetic field. This design is similar to simulated annealing of an Ising Hamiltonian, with the key difference being that the annealing process avoids getting trapped in local minima via quantum tunneling instead of thermal fluctuations, and the probability of hopping out of a local minimum is determined by the width of the barrier rather than the height (35). AQCs therefore provide a compelling strategy for solving combinatorial optimization problems that can be broken down into one- and two-body interactions, and these types of problems could offer quantum advantage for practical use-cases in the near term (36). For example, a recent study explored the potential for leveraging AQCs and gate-based devices for codon optimization, a crucial process in reagent generation and mRNA vaccine development (37).

In this study we show that the RNA secondary structure prediction problem can be mathematically formulated as a BQM and thus be addressed using quantum computing technology. This representation requires translating the objective function utilized in classical approaches to a polynomial Hamiltonian where the eigenstates represent combinations of secondary structure elements, and the eigenvalues represent the scores. Implementing this approach on the D-Wave Advantage 1.1 hybrid solver provides performance competitive with a parallel replica exchange Monte Carlo (REMC) algorithm utilizing 128 cores programmed with the same objective function.

## Results

RNA secondary structure prediction was implemented as a BQM on a D-Wave AQC with a Hamiltonian designed to optimize the number of non-overlapping, consecutive, intramolecular base pairs with penalties imposed for pseudoknots and additional constraints to prevent bases from forming more than one hydrogen bond. The list of all possible stems is pre-computed classically using the methods described in (24). Figure 1a shows an example matrix formulation for sequence: GGAAGCAAACAUCCCUGU, and Figure 1b provides a visualization of the base pairing patterns identified in Figure 1a. The classical data was encoded onto the quantum device by mapping each stem to a qubit (Figure 1c). The qubits which return “1” upon measurement represent the stems contributing to the secondary structure. The final secondary structure pattern is determined by recording the values of the qubits and saving all stems represented by qubits that returned “1”.

**Figure 1:**
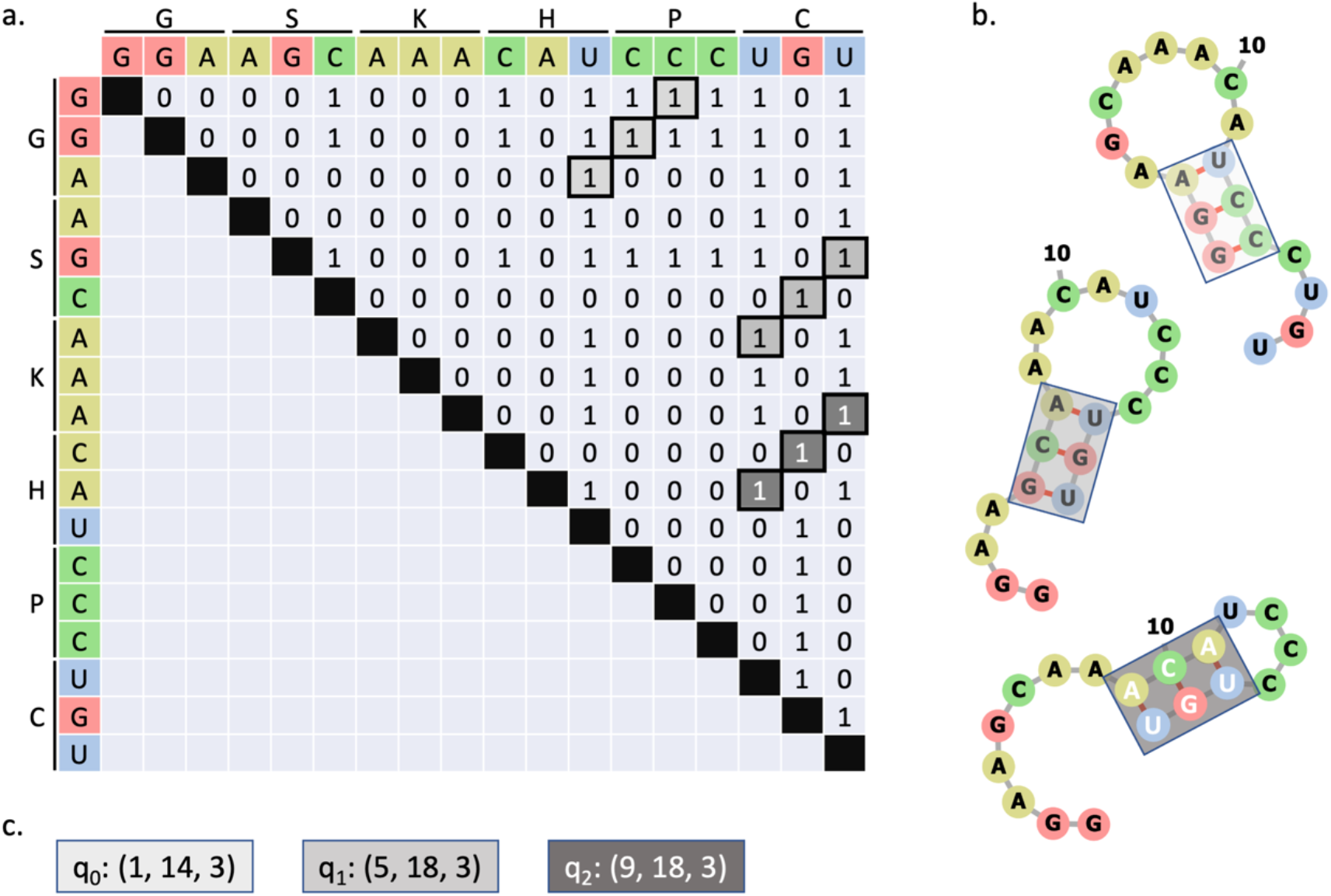
(a) Matrix representation of potential hydrogen bonds. Stems are highlighted in orange, blue, and green. (b) Structural representations of stems. Grey lines indicate covalent bonds and red lines indicate intramolecular hydrogen bonds. The color of the boxes map to the stems identified in (a). (c) Each stem is assigned to a qubit on the quantum device. If the qubit returns “1”, then the associated stem is included in the RNA secondary structure. Otherwise, the stem is excluded.

Directly programming the Quantum Processing Unit (QPU) with the BQM yields noisy data. Supplementary Figure 1 shows the distribution of scores for systems requiring up to 45 physical qubits. There are a multitude of factors that contribute to the noise, such as thermal fluctuations, and in general the noise is exacerbated by increasing the number of qubits. However, D-Wave offers a hybrid solver which uses a classical device to break down the problem into smaller pieces that are handled by the QPU. The following sections compare the performance of the hybrid solver against a REMC algorithm programmed with the same objective function and exact enumeration of the solution space where possible.

### Exact Enumeration (<= 45 stems)

Problems with sufficiently small solution spaces can be solved exactly. An MPI-enabled solver was written (see Methods) to exhaustively solve systems using massively parallel resources. Using this method, systems containing up to 45 qubits (2^45^ (~10^13^) possible solutions) can be solved in under 24 hours using 7,900 CPU cores. Comparing the lowest energy solution to the experimentally determined solution (referred to as the known solution) is used to assess the suitability of the objective function. The structure predicted by the algorithm is compared to the known solution by computing the sensitivity and the specificity. The sensitivity reflects the fraction of experimentally determined base pairs that were correctly identified. A high sensitivity score (maximum score is 1) means the algorithm predicted every naturally occurring base pair and a low sensitivity score (minimum score is 0) means the algorithm did not predict many of the naturally occurring base pairs. The specificity reflects the fraction of predicted base pairs that map to the naturally occurring structure. A specificity score of 1 indicates that the algorithm only predicted base pairs that match the known structure, and a low score indicates that the algorithm predicted many base pairs that do not map to the known structure.

A test set was derived by scraping PseudoBase for examples of RNA sequences with experimentally confirmed structures containing pseudoknots (38–40). Redundant sequences, defined as sequences with higher than 95% similarity, were removed from the set. The number of possible stems was computed for the remaining sequences in the database, and it was found that most sequences yielded too many possible stems for exact enumeration. To reduce the total number of possible stems down to a set small enough for exact enumeration, the length of the smallest possible stem was increased to 4. This restriction excludes stems of lengths 2 and 3 from the sets, thereby reducing the size of the combinatorial solution space. Imposing this constraint resulted in a test set containing 27 sequences. Figure 2a shows an example pseudoknotted structure from the test set. Figure 2b shows the sensitivity of the objective function was calculated to be 0.97 and the specificity was found to be 0.93, indicating that 97% of the naturally occurring base pairs were recapitulated by the algorithm, and 93% of the base pairs predicted by the algorithm correctly map to the naturally occurring structures. The full list of results can be found in Supplementary Table 1.

**Figure 2.**
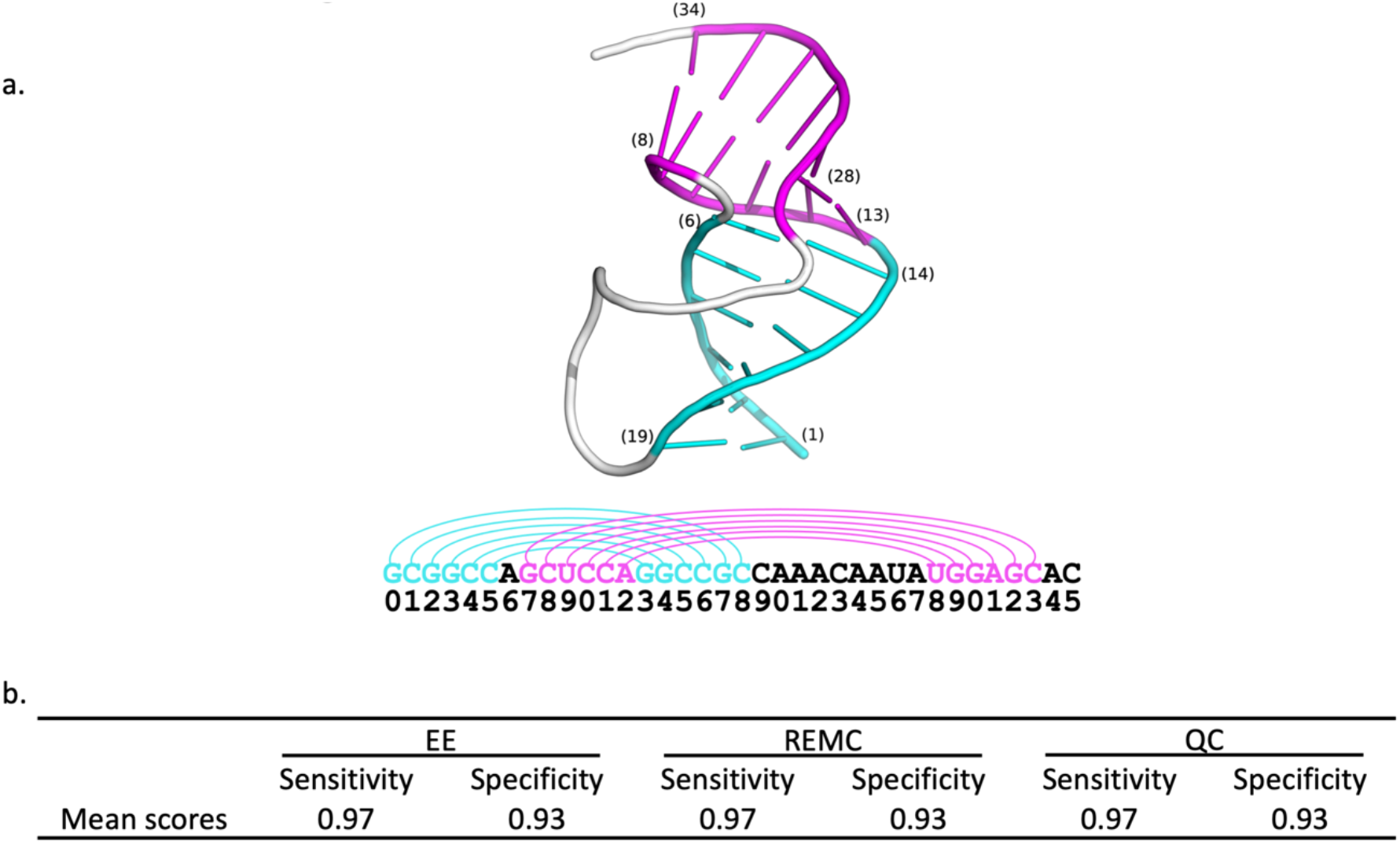
(a) Example RNA sequence from simian retrovirus type-1 (SRV-1) with structure containing pseudoknot correctly predicted by algorithm (PDB code: 1E95). The phosphate backbone is colored white for unpaired bases. (b) Exact Enumeration (EE) was used to compute the sensitivity and specificity of the proposed Hamiltonian (Equation 12) for systems containing 45 stems or fewer. Replica Exchange Monte Carlo (REMC) and quantum computing (QC) methods were tested on the same sequences and were found to exactly recapitulate the lowest energy solutions found by exact enumeration.

The same analysis was performed using the REMC and QC methods. The average results are shown in Figure 2b, and the full list of results are shown in Supplementary Table 1. Each system was run 1 time through each algorithm, and in every case the result obtained matched the result from exact enumeration. Therefore, both methods are able to rapidly identify the minimum energy solution with high probability for systems containing fewer than 45 stems.

### Simulated annealing vs. quantum computing (>45 stems)

Scaling the system size beyond 45 stems requires tremendous computational resources for exact enumeration therefore these systems are only evaluated against approximate methods. Systems containing 45 – 881 stems were evaluated using both the REMC and QC methods.

Similar to the previous section, the test sequences derive from PseudoBase. However, the minimum stem length was set to 3 and the minimum loop length was set to 2, allowing for significantly larger problem sizes. Each sequence was run through both algorithms 10 times to estimate the variance of the results. The sensitivity and specificity in these cases are less indicative of the fitness of the objective function since it is unlikely that the true minimum energy solution will be found each time. Figure 3a compares the scores of the REMC and QC methods. Points that fall on the dashed line indicate that the same exact score was found for both methods. Points that fall below the line indicate that the QC method found a lower energy solution, and similarly points above the line indicate the REMC method found a better solution. On average, the methods produce results of similar quality. Figure 3b shows the ensemble average sensitivity and specificity for each method. The REMC method reported slightly higher sensitivity and specificity than the QC method.

**Figure 3.**
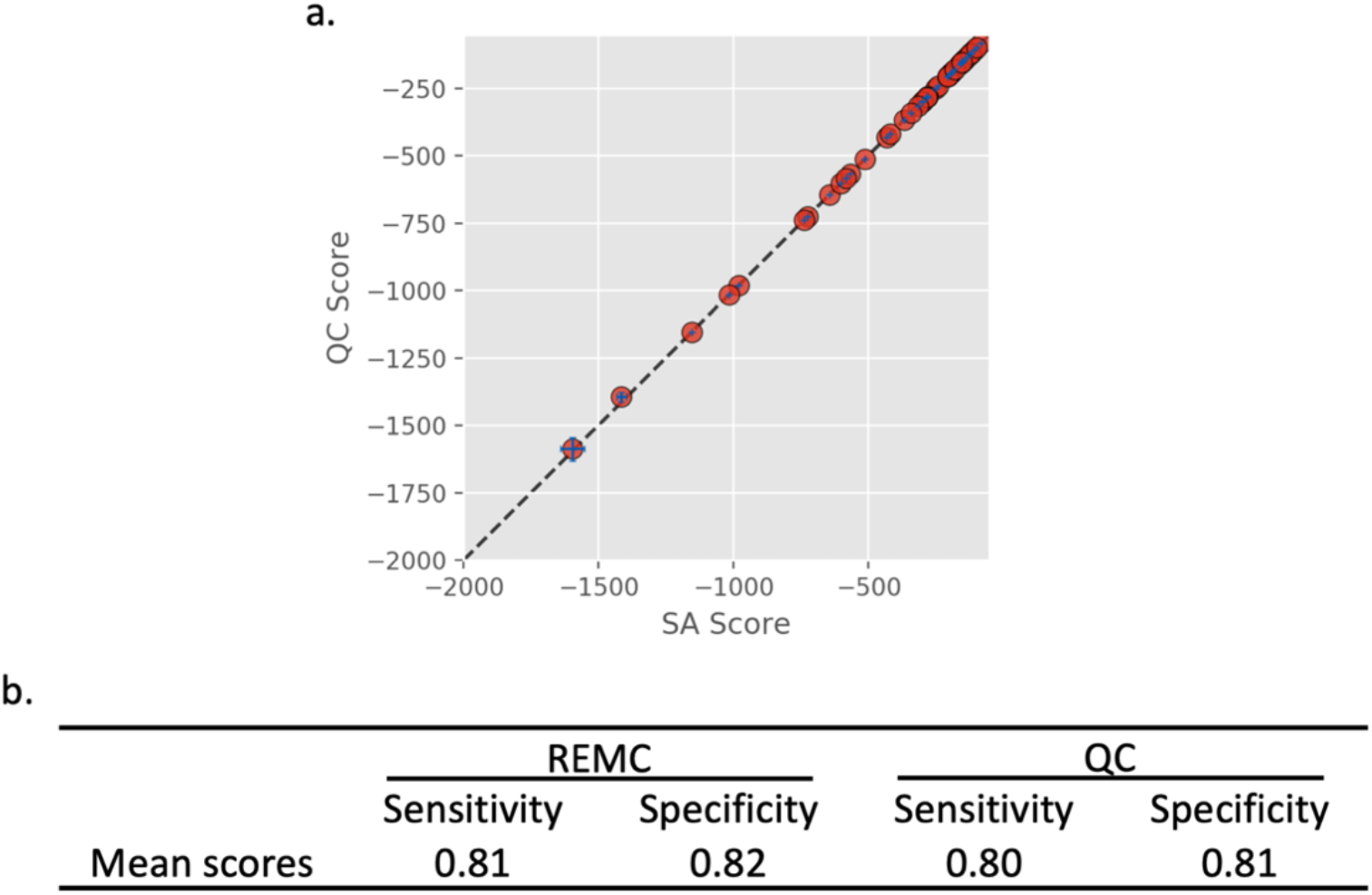
Replica Exchange Monte Carlo (REMC) and Quantum Computing algorithms were tested on systems containing more than 45 stems, which exceeds the practical limit of what can be solved with exact enumeration. The eigenvalues of the lowest energy systems are compared in (a). The dashed line represents y=x.

### Comparison to existing methods

The results presented in the previous sections were compared to three algorithms found in the literature. Two of the methods, SPOT-RNA (11) and ProbKnot (41), are capable of predicting pseudoknots while the other method, ViennaRNA (42), is not. The algorithms were tested on the same datasets as the previous sections. Figure 4 shows a summary of the overall sensitivity and specificity for each method applied to each dataset, listed in descending order. The method presented in this study had the best overall performance on both datasets. All methods performed worse on the dataset containing larger systems (>=45 stems) compared to the dataset containing smaller systems (<45 stems).

**Figure 4.**
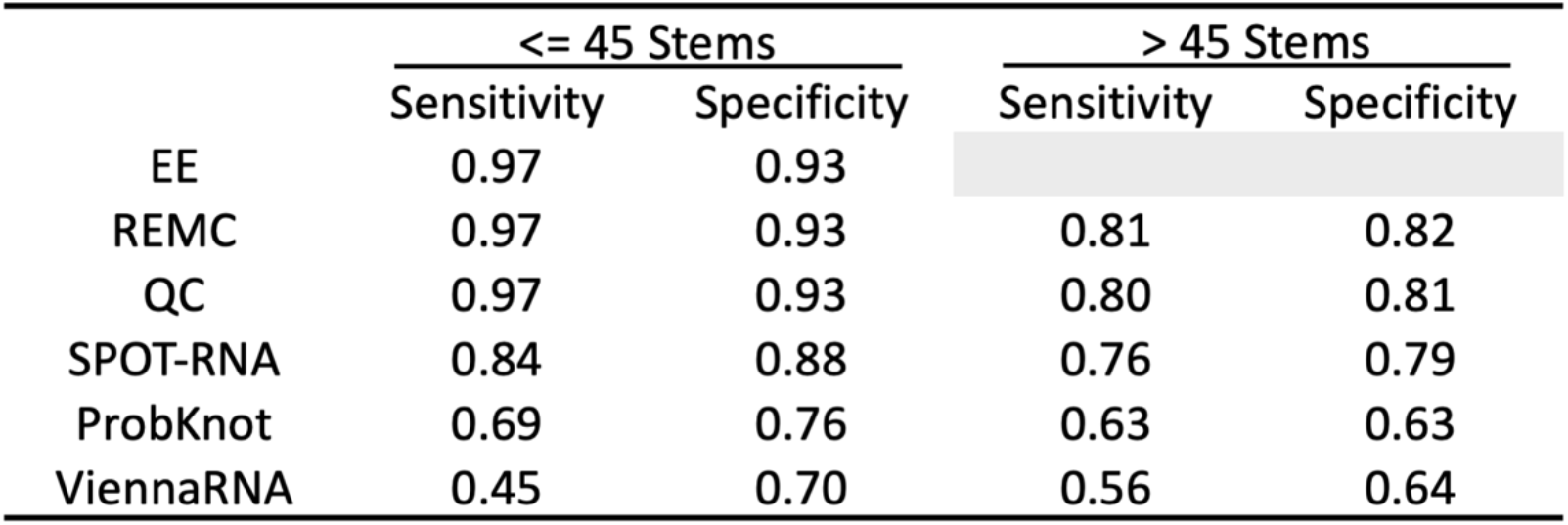
Comparison of methods on the datasets described in previous sections. Exact Enumeration (EE) is not computed for systems containing more than 45 stems.

ViennaRNA yielded the poorest agreement with both datasets. Given that the method is not advertised to predict pseudoknots, this was the expected outcome. SPOT-RNA, a recently proposed method using an ensemble of deep neural networks, performed almost as well as the QC on the dataset containing sequences with more than 45 stems and had overall the most consistent performance between the two datasets.

## Discussion

The purpose of this study was to present a model that enables RNA structure prediction using existing quantum hardware. AQCs were chosen to demonstrate the viability of the method due to the relative maturity of the technology, but BQM’s are readily implementable on all types of QCs. A recent study compared the performance and accuracy of AQCs and gate-based models for a BQM describing a biological system and showed that the gate-based platforms lack the qubits and physical connectivity required to test realistically large systems (37). However, there are proposals to build gate-based error corrected devices with up to 1,000,000 qubits by 2025 (43). Such a device would be capable of encoding the 10^146,440^ possible solutions to the RNA structure of the SARS-CoV2 spike glycoprotein. The ability to find better approximations to massive combinatorial problems like this could have a tremendous impact on vaccine design and drug discovery.

The objective function used in this study is designed to simultaneously maximize the number of base pairs and the average length of the stems. There are cases where the sampling methods identify patterns of base pairs that score higher in our metric than what is observed in nature, indicating that the scoring function does not perfectly recapitulate the physics driving RNA folding. However, despite the simplicity of the scoring function, it was shown that the algorithm correctly identified the natural base pairs with 97% sensitivity and 93% specificity in smaller cases where exact enumeration was possible. There were 4 cases where either the sensitivity or the specificity was below 0.75. In one of the cases, the algorithm preferred a configuration containing four stems of length five instead of two stems, one of length five and one of length six. In the other 3 cases the algorithm found solutions containing more base pairs with at least 70% specificity.

In larger cases where exact enumeration is impossible, both the quantum and the REMC methods yielded results of similar quality. In general, larger systems tended to have poorer agreement with naturally occurring structures. Overall, the method presented in this study outperformed the three comparator methods in terms of sensitivity and specificity. However, the datasets used in this study only included structures containing pseudoknots, but there are many classes of RNAs and many other types of secondary structure that need to be tested for a more thorough understanding of where the method is applicable.

The objective function used in this study was designed to be simple and interpretable, but there is room for improvement. For example, the force field could take into account the number of hydrogen bonds formed by each type of interaction which could potentially reduce the number of degenerate low-lying states. Furthermore, base pairs involved in pseudoknots are penalized by scaling back their energetic contribution to the Hamiltonian. The energy contribution for such base pairs is expected to be smaller because the formation of the knot requires the backbone to bend which puts stress on the hydrogen bonded base pairs. For simplicity, this effect was accounted for as a constant, but a systematic structural study could be performed to derive a more realistic model.

It should be noted that the purpose of this work was not to develop a higher fidelity RNA folding potential but rather to show how one could develop a classical folding potential that can be easily mapped to quantum hardware. As long as one sticks to a discrete polynomial formalism more complex potentials can be mapped to existing and future QCs with ease. Hence in conclusion QCs are very promising for RNA folding. They are able to rapidly find minimum energy solutions for large combinatorial search spaces and have the flexibility to employ more complex potentials which should deliver higher specificity and sensitivity with limited impact on computational time. As the capabilities of QCs improve, in terms of qubit count, connectivity and signal to noise ratios it will become feasible to fold very large RNA sequences that would be intractable using classical methods.

## Methods

### RNA Secondary Structure Prediction Algorithm

For a given RNA sequence containing *N* bases, an *NxN* matrix in constructed where rows and columns represent the bases. The upper diagonal elements of the matrix are populated with 1’s for combinations of bases that hydrogen bond each other (A-U, G-C, G-U) and 0’s for combinations that do not form hydrogen bonds. For a reasonably strong interaction to persist in an RNA structure, a minimum of 3 consecutive hydrogen bonds are required. The simplest way to identify consecutive bonds, called *stems*, is to scan the matrix for repeated 1’s in a diagonal perpendicular to the diagonal of the matrix. Figure 1a shows the matrix construction and corresponding hydrogen bonding patterns for an example sequence, GGAAGCAAACAUCCCUGU. Consecutive hydrogen bonds (with three or more bonds in a row) are highlighted in varying shades of gray, and representations of RNA folds subjected to these bonding patterns are displayed in Figure 1b.

### Implementation of objective function

The goal of the optimization algorithm is to identify the combination of non-overlapping stems that simultaneously maximizes the number of consecutive base pairs along with the average length of the chosen set of stems. There are three parts to the global objective function; the first term reflects the number of base pairs with an adjustable penalty accounting for pseudoknots, the second term adds a penalty for adding short stems, and finally a constraint which adds an infinite penalty to combinations of overlapping stems.

### Scoring consecutive base pairs

The base pair scoring function is inspired by the fitness function from Kai et al (24) which takes into consideration the number of consecutive base pairs and the number of pseudoknots. While there are many approaches to scoring base pairing networks, this approach was chosen due to its simplicity. Each stem is parameterized by the index of the first base, *i*, the index of the last base, *j*, and the length of the stem, *k*. This is compactly written as *m*_*n*_ = (*i*_*n*_, *j*_*n*_, *k*_*n*_) for stem *n*. For a set of stems *M* = {*m*_*0*_, *m*_*1*_, …, *m*_*N*_}, the average number of consecutive bonds, *b*, is computed by summing over the lengths of each stem divided by the number of stems:

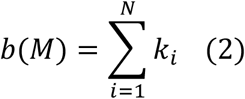

To prioritize base pairing configurations with longer stems, the Hamiltonian incorporates Equation (2) squared.

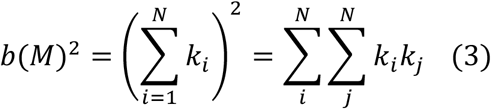

Equation (3) can be formulated as the Hamiltonian of a quantum system by taking the inner product with the vector of qubits comprising the quantum state, q = (*q*_*0*_, *q*_*1*_, …, *q*_*N*_), where *q*_*i*_ ∈ {0,1}, and *c*_*B*_ is a tunable constant:

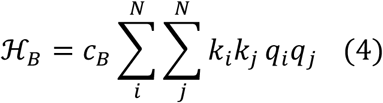

This representation maps each stem to a labeled qubit, and the values of the binary values of the qubits determine which stems from the set contribute to the overall sum.

The matrix represented in the double sum needs to be restricted to a sum over the upper triangular elements, consistent with Equation (1). By decomposing the sum into the trace and a term that sums the contributions of the off-diagonal elements, the sum can be restricted to the upper triangular elements. The trace requires a single summation over *k*_*i*_^*2*^. Since qubits map to binary values, they are idempotent with themselves and therefore *q*_*i*_^*2*^ = *q*_*i*_.

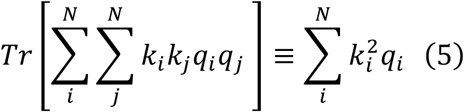

Since the matrix is symmetric, all off-diagonal terms are accounted for in an upper triangular form by multiplying by 2. Thus,

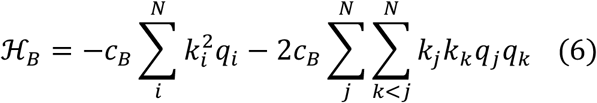

### Incorporating pseudoknot penalties

A pseudoknot is defined as two non-overlapping stems, *m*_*a*_ = (*i*_*a*_, *j*_*a*_, *k*_*a*_) and *m*_*b*_ = (*i*_*b*_, *j*_*b*_, *k*_*b*_) where *i*_*a*_ < *i*_*b*_ < *j*_*a*_ < *j*_*b*_ or *i*_*b*_ < *i*_*a*_ < *j*_*b*_ < *j*_*a*_. Pseudoknots are detected by a delta function,

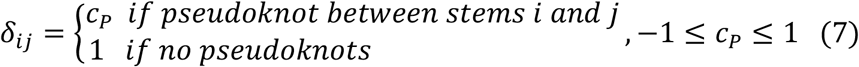

Where *c*_*P*_ is a tunable parameter set by the user. This penalty is designed to reduce the contribution of hydrogen bonds from pseudoknots. When *c*_*P*_ is set to 1, hydrogen bonds from pseudoknots are not penalized. Conversely, when *c*_*P*_ is set below 0, pseudoknots will not exist in the global minimum and can be avoided altogether. If *c*_*P*_ is set to zero, the global minimum could be degenerate with structures containing pseudoknots and the same structures with the pseudoknots removed. Incorporating this factor into the Hamiltonian yields Equation (8).

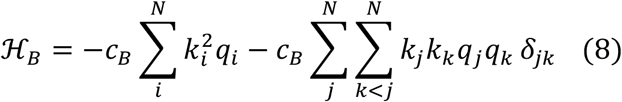

The delta-function is omitted from the single summation since a stem cannot be in a pseudoknot with itself, and therefore the function will always evaluate as 1.

### Maximize average length of stem

One of the distinguishing features of the objective function introduced in (24) is an energetic preference for longer stems. Longer stems are assigned lower energetic values in Equation (8), but this term does not distinguish between multiple short stems and one long stem. For example, two stems of length three would yield a score of (3+3)^2^ = 36, whereas a single stem of length six would yield a score of 6^2^ = 36. A single stem of length 6 would be preferred in most cases to two separate stems of lengths three because a penalty is incurred for interrupting the contiguous pi-pi interactions between bases. Since there are exceptions (see Results), this feature is incorporated into the Hamiltonian as a tunable parameter c_L_.

The average stem length can be maximized by minimizing the difference between the length of each stem in the set, *k*_*i*_, with the length of the largest stem in the set, *μ*.

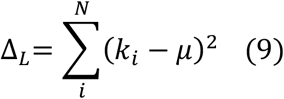

Equation 9 is rewritten as a Hamiltonian by expanding the product and projecting with the vector representing the qubits, **q**.

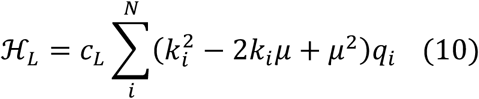

### Hamiltonian with constraints

Bases are only able to form hydrogen bonds with exactly one other base, so combinations of stems that require more than one bond for any given base must be excluded from the solution space. A delta-function is introduced to detect such combinations:

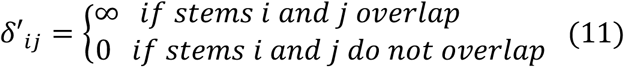

The total Hamiltonian is thus constructed by adding the Hamiltonian defined in Equation (8) with the constraint defined in Equation (11):

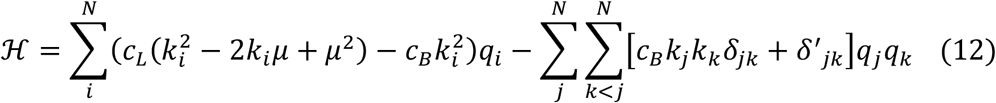

### Algorithm implementations

The current approach to performing calculations on quantum devices requires the interaction terms to be precomputed on classical devices and read into the quantum devices via specialized APIs. The interaction parameters from Equation (12) were precomputed in python 3.7 using standard libraries and numpy arrays (44). The numpy arrays were converted to dictionaries in accordance with the expected input for the D-Wave libraries. The execution of the BQM was carried out using libraries described in the following sections. Each calculation was run 10 times. The pseudoknot penalty, *c*_*P*_, was set to 0.5 for all calculations.

### D-Wave Advantage 1.1

The RNA secondary structure BQM was implemented on the D-Wave Advantage System 1.1 utilizing the Leap Hybrid Solver. This adiabatic quantum device contains more than 5,000 superconducting qubits. Each qubit is connected to 15 others described by a Pegasus P_16_ graph (45). The Advantage system was accessed through the D-Wave Leap web interface, which serves as an access point to QPU hardware as well as an integrated developer environment with built-in support for the full D-Wave API.

The program was constructed and executed using python libraries provided by D-Wave systems. The BinaryQuadraticModel class in the dimod 0.9.10 python library was used to construct the model from the classically prepared data and convert it to a data structure compatible with the quantum device. The one- and two-body interaction terms were precomputed, stored in numpy arrays, and passed into the BinaryQuadraticModel instance along with an offset of 0.0 as a dimod.BINARY representation. The model was executed using the LeapHybridSampler classes in the dwave.system python library. The solver was allotted 3 s of execution time. The eigenstate with the lowest associated eigenvalue was chosen to represent the result of the simulation.

### Replica Exchange Monte Carlo

Replica exchange Monte Carlo (REMC) is a well-established and widely used method for identifying global minima on high complexity objective surfaces that contain many local minima (46–48). Here, a very basic REMC search algorithm was implemented to explore the folding landscape with the following steps: 1. An initial state is created composed of up to 5 randomly selected stems from the set of all stems. 2. The objective function is evaluated for this initial state. 3. A Monte Carlo move is performed in which three changes to the state are possible, each with equal probability: a) a stem is added, b) a stem is removed, or c) a stem is swapped for another from the set of all stems. In each case, if the move improves the objective function, the state is accepted, otherwise the state is accepted subject to the Metropolis-Hastings criteria (49) with a probability proportional to the Boltzmann distribution at a particular temperature. Subsequent Monte Carlo moves are then performed a fixed number of times, after which the final state of the system is returned. The Monte Carlo search when combined with simulated annealing and exponential cooling was found to be effective in identifying the global minimum of the objective for systems having fewer than 45 stems. However, as the complexity of the objective surface grows with increasing numbers of stems and possible states, obtaining effective sampling of the landscape becomes challenging with simulated annealing, often requiring increasingly complex cooling schedules, continuous tuning of the initial sampling temperature, or drastically increasing the number of sampling steps (50).

The MC method was designed to allow for the simultaneous evolution of *N* replicas, each at a fixed effective temperature. Replicas sample states with probabilities proportional to the Boltzmann distribution at that replica’s temperature. The stochastic nature of the algorithm and the initial random configuration of the system necessitates that the replicas be able to exchange states with those at neighboring temperatures. Without exchange, replicas at lower temperatures may become stuck in the numerous local minima of the objective function, while those at higher temperatures will never sample the desired higher probability states. Thus, by allowing swapping of states of neighboring temperatures, lower probability states are sampled in replicas at higher temperatures, which are then exchanged with replicas at lower temperatures to sample the desired higher probability states (46–48).

In this implementation, exchanging states between replicas is performed according to the Metropolis-Hastings criteria (49), with a probability proportional to the difference in the objective’s value between the two states and their temperatures. Exchange attempts are performed every fixed number of steps, allowing for independent MC sampling to occur in each replica between exchange attempts, ensuring detailed balance. The replica temperature range, as well as their spacing are selected such that exchanges between neighboring replicas occurs roughly twenty percent of the time. The final state of the system is chosen from the best scoring state from the collection of replicas following the final step.

Using this REMC approach, hundreds of potential stems could be efficiently sampled using only 64 replicas, with values of β (1/kT, where k is the Boltzmann constant and T is the effective temperature) in arbitrary units ranging from 1 to 30, 10^6^ total steps, and exchange attempts every 10^3^ steps. The total time to identify the global minimum for a system composed of 400 stems is roughly twelve seconds.

The REMC method was implemented in pure Python and takes advantage of NumPy (44), and Python’s Random and Copy libraries. Replica parallelization, including synchronization and swapping of replica states was implemented using MPI for Python (51–53).

### Comparator methods

Three additional, previously published methods were run using the same datasets and the same criteria for comparing to known structures as the method proposed in this study. RNAstructure ProbKnot 6.3 (41), SPOT-RNA (11), and ViennaRNA RNAfold 2.4.12 (42) were all run locally on an HPC cluster using command line defaults. The iterations parameter required by ProbKnot was set to 1000. The base pairs were extracted by mining the output PostScript (ps) file (ViennaRNA) or Connectivity Table (ct) file (ProbKnot and SPOT-RNA).

### Metrics for comparing stems

The predicted secondary structure was compared to the experimentally determined structure by computing the sensitivity and specificity. The sensitivity, *σ* _*SN*_, is computed by comparing the number of correctly identified base pairs, *C*, to the number of predicted base pairs missing from the known structure, *M*, with the following formula:

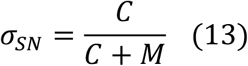

The specificity, *σ*_*SP*_, is computed by comparing *C* to the number of predicted base pairs that are not represented in the known structure, *I*.

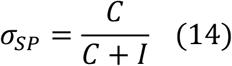

## Acknowledgements

We would like to thank Kim Branson, Deborah Loughney, Laura Zupko, and Lee Tremblay of GSK for evaluation and suggested refinements.

## Competing Interests

The authors declare no competing interests.

## Notes

### Competing Interest Statement

The authors have declared no competing interest.

## References

1. Cooper GM. The Cell: A Molecular Approach. 2nd edition [Internet]. Sinauer Associates 2000; 2000. Available from: http://lib.ugent.be/catalog/ebk01:3450000000002155

2. Amaral PP, Dinger ME, Mercer TR, Mattick JS. The eukaryotic genome as an RNA machine. Science (80-). 2008;319(5871):1787–9.

3. Serganov A, Patel DJ. Ribozymes, riboswitches and beyond: Regulation of gene expression without proteins. Nat Rev Genet. 2007;8(10):776–90.

4. Chemla Y, Peeri M, Heltberg ML, Eichler J, Jensen MH, Tuller T, et al. A possible universal role for mRNA secondary structure in bacterial translation revealed using a synthetic operon. Nat Commun [Internet]. 2020;11(1):1–11. Available from: http://dx.doi.org/10.1038/s41467-020-18577-4

5. Gorochowski TE, Ignatova Z, Bovenberg RAL, Roubos JA. Trade-offs between tRNA abundance and mRNA secondary structure support smoothing of translation elongation rate. Nucleic Acids Res. 2015;43(6):3022–32.

6. Cambray G, Guimaraes JC, Arkin AP. Evaluation of 244,000 synthetic sequences reveals design principles to optimize translation in escherichia coli. Nat Biotechnol. 2018;36(10):1005.

7. Bellaousov S, Reuter JS, Seetin MG, Mathews DH. RNAstructure: Web servers for RNA secondary structure prediction and analysis. Nucleic Acids Res. 2013;41(Web Server issue).

8. Andronescu M, Condon A, Hoos HH, Mathews DH, Murphy KP. Efficient parameter estimation for RNA secondary structure prediction. Bioinformatics. 2007;23(13):19–28.

9. Zuker M, Stiegler P. Optimal computer folding of large RNA sequences using thermodynamics and auxiliary information. Nucleic Acids Res. 1981;9(1):133–48.

10. Senior AW, Evans R, Jumper J, Kirkpatrick J, Sifre L, Green T, et al. Improved protein structure prediction using potentials from deep learning. Nature [Internet]. 2020;577(7792):706–10. Available from: http://dx.doi.org/10.1038/s41586-019-1923-7

11. Singh J, Hanson J, Paliwal K, Zhou Y. RNA secondary structure prediction using an ensemble of two-dimensional deep neural networks and transfer learning. Nat Commun [Internet]. 2019;10(1). Available from: http://dx.doi.org/10.1038/s41467-019-13395-9

12. Zhang H, Zhang C, Li Z, Li C, Wei X, Zhang B, et al. A new method of RNA secondary structure prediction based on convolutional neural network and dynamic programming. Front Genet. 2019;10(MAY):1–12.

13. Lu W, Tang Y, Wu H, Huang H, Fu Q, Qiu J, et al. Predicting RNA secondary structure via adaptive deep recurrent neural networks with energy-based filter. BMC Bioinformatics [Internet]. 2019;20(Suppl 25):1–10. Available from: http://dx.doi.org/10.1186/s12859-019-3258-7

14. Fallmann J, Will S, Engelhardt J, Grüning B, Backofen R, Stadler PF. Recent advances in RNA folding. J Biotechnol [Internet]. 2017;261(July):97–104. Available from: http://dx.doi.org/10.1016/j.jbiotec.2017.07.007

15. Bringmann K, Grandoni F, Saha B, Williams VV. Truly subcubic algorithms for language edit distance and RNA folding via fast bounded-difference min-plus product. SIAM J Comput. 2019;48(2):481–512.

16. Huang L, Zhang H, Deng D, Zhao K, Liu K, Hendrix DA, et al. LinearFold: Linear-time approximate RNA folding by 5’-to-3’ dynamic programming and beam search. Bioinformatics. 2019;35(14):i295–304.

17. Lyngsø RB, Pedersen CNS. RNA pseudoknot prediction in energy-based models. J Comput Biol. 2000;7(3–4):409–27.

18. Messenger RNA encoding the full-length SARS-CoV-2 spike glycoprotein [Internet]. WHO MedNet; 2020. Available from: https://web.archive.org/web/20210105162941/https://mednet-communities.net/inn/db/media/docs/11889.doc

19. Alves J, Combes F, Ferrara A, Forveille T, Shore S. Planck 2015 results. Astron Astrophys. 2016;594.

20. Hofacker IL. Vienna RNA secondary structure server. Nucleic Acids Res. 2003;31(13):3429–31.

21. Mathews DH. Revolutions in RNA Secondary Structure Prediction. J Mol Biol. 2006;359(3):526–32.

22. Knudsen B, Hein J. Pfold: RNA secondary structure prediction using stochastic context-free grammars. Nucleic Acids Res. 2003;31(13):3423–8.

23. Montaseri S, Zare-Mirakabad F, Moghadam-Charkari N. RNA-RNA interaction prediction using genetic algorithm. Algorithms Mol Biol. 2014;9(1):1–7.

24. Kai Z, Yuting W, Yulin L, Jun L, Juanjuan H. An efficient simulated annealing algorithm for the RNA secondary structure prediction with Pseudoknots.BMC Genomics [Internet]. 2019;20(Suppl 13):1–13. Available from: http://dx.doi.org/10.1186/s12864-019-6300-2

25. Zhong H-S, Wang H, Deng Y-H, Chen M-C, Peng L-C, Luo Y-H, et al. Quantum computational advantage using photons. Science (80-) [Internet]. 2020;1463(December):1460–3. Available from: http://science.sciencemag.org/

26. Arute F, Arya K, Babbush R, Bacon D, Bardin JC, Barends R, et al. Quantum supremacy using a programmable superconducting processor. Nature [Internet]. 2019;574(7779):505–10. Available from: http://dx.doi.org/10.1038/s41586-019-1666-5

27. Bova F, Goldfarb A, Melko RG. Commercial applications of quantum computing. EPJ Quantum Technol [Internet]. 2021;8(1):1–13. Available from: http://dx.doi.org/10.1140/epjqt/s40507-021-00091-1

28. Cheng HP, Deumens E, Freericks JK, Li C, Sanders BA. Application of Quantum Computing to Biochemical Systems: A Look to the Future. Front Chem. 2020;8(November):1–13.

29. Elfving VE, Broer BW, Webber M, Gavartin J, Halls MD, Lorton KP, et al. How will quantum computers provide an industrially relevant computational advantage in quantum chemistry? arXiv. 2020;1–20.

30. Wang H, Kais S, Aspuru-Guzik A, Hoffmann MR. Quantum algorithm for obtaining the energy spectrum of molecular systems. Phys Chem Chem Phys. 2008;10(35):5388–93.

31. Aspuru-Guzik A, Dutoi AD, Love PJ, Head-Gordon M. Chemistry: Simulated quantum computation of molecular energies. Science (80-). 2005;309(5741):1704–7.

32. Cao Y, Romero J, Olson JP, Degroote M, Johnson PD, Kieferová M, et al. Quantum Chemistry in the Age of Quantum Computing. Chem Rev. 2019;119(19):10856–915.

33. Kassal I, Jordan SP, Love PJ, Mohseni M, Aspuru-Guzik A. Polynomial-time quantum algorithm for the simulation of chemical dynamics. Proc Natl Acad Sci U S A. 2008;105(48):18681–6.

34. Kühn M, Zanker S, Deglmann P, Marthaler M, Weiß H. Accuracy and Resource Estimations for Quantum Chemistry on a Near-Term Quantum Computer. J Chem Theory Comput. 2019;15(9):4764–80.

35. Muthukrishnan S, Albash T, Lidar DA. Tunneling and speedup in quantum optimization for permutation-symmetric problems. Phys Rev X. 2016;6(3):1–23.

36. Djidjev H, Chapuis G, Hahn G, Rizk G. Efficient combinatorial optimization using quantum annealing. arXiv. 2018;1–25.

37. Fox DM, Branson KM, Walker RC. mRNA codon optimization on quantum computers. bioRxiv [Internet]. 2021;2021.02.19.431999. Available from: https://doi.org/10.1101/2021.02.19.431999

38. Van Batenburg FHD, Gultyaev AP, Pleij CWA, Ng J, Oliehoek J. PseudoBase: A database with RNA pseudoknots. Nucleic Acids Res. 2000;28(1):201–4.

39. Van Batenburg FHD, Gultyaev AP, Pleij CWA. PseudoBase: Structural information on RNA pseudoknots. Nucleic Acids Res. 2001;29(1):194–5.

40. Taufer M, Licon A, Araiza R, Mireles D, van Batenburg FHD, Gultyaev AP, et al. PseudoBase++: An extension of PseudoBase for easy searching, formatting and visualization of pseudoknots. Nucleic Acids Res. 2009;37(SUPPL. 1):127–35.

41. Bellaousov S, Mathews DH. ProbKnot: Fast prediction of RNA secondary structure including pseudoknots. Rna. 2010;16(10):1870–80.

42. Lorenz R, Bernhart SH, Höner zu Siederdissen C, Tafer H, Flamm C, Stadler PF, et al. ViennaRNA Package 2.0. Algorithms Mol Biol. 2011;6(1):1–14.

43. Smith-Goodson P. Quantum Computing With Particles Of Light: A $ 215 Million Gamble. Forbes [Internet]. 2020 Apr; Available from: https://www.forbes.com/sites/moorinsights/2020/04/15/quantum-computing-with-particles-of-light-a-215-million-gambl/?sh=13b4fb5224a7

44. Harris CR, Millman KJ, van der Walt SJ, Gommers R, Virtanen P, Cournapeau D, et al. Array programming with {NumPy}. Nature [Internet]. 2020 Sep;585(7825):357–62. Available from: https://doi.org/10.1038/s41586-020-2649-2

45. Dattani N, Szalay S, Chancellor N. Pegasus: The second connectivity graph for large-scale quantum annealing hardware. arXiv. 2019;

46. Geyer CJ, Thompson EA. Annealing Markov Chain Monte Carlo with Applications to Ancestral Inference. J Am Stat Assoc [Internet]. 1995;90(431):909–20. Available from: https://www.tandfonline.com/doi/abs/10.1080/01621459.1995.10476590

47. Koji H, Koji N. Exchange Monte Carlo Method and Application to Spin Glass Simulations. J Phys Soc Japan [Internet]. 1996;65(6):1604–8. Available from: https://doi.org/10.1143/JPSJ.65.1604

48. Swendsen RH, Wang J-S. Replica Monte Carlo Simulation of Spin-Glasses. Phys Rev Lett [Internet]. 1986 Nov 24;57(21):2607–9. Available from: https://link.aps.org/doi/10.1103/PhysRevLett.57.2607

49. Hastings WK. Monte Carlo sampling methods using Markov chains and their applications. Biometrika [Internet]. 1970 Apr 1;57(1):97–109. Available from: https://doi.org/10.1093/biomet/57.1.97

50. Kofke DA. On the acceptance probability of replica-exchange Monte Carlo trials. J Chem Phys [Internet]. 2002 Sep 30;117(15):6911–4. Available from: https://doi.org/10.1063/1.1507776

51. Dalcín L, Paz R, Storti M. MPI for Python. J Parallel Distrib Comput. 2005;65(9):1108–15.

52. Dalcin LD, Paz RR, Kler PA, Cosimo A. Parallel distributed computing using Python. Adv Water Resour [Internet]. 2011;34(9):1124–39. Available from: http://dx.doi.org/10.1016/j.advwatres.2011.04.013

53. Dalcín L, Paz R, Storti M, D’Elía J. MPI for Python: Performance improvements and MPI-2 extensions. J Parallel Distrib Comput. 2008;68(5):655–62.

