## Supplemental Information for "RNA folding using quantum computers"

### Supplementary Information

#### Scores from individual sequences

| Name | EMBL Number | EE | | REMC | | QC | |
|---|---|---|---|---|---|---|---|
| | | Sensitivity | Specificity | Sensitivity | Specificity | Sensitivity | Specificity |
| FIV | M25381 | 1.00 | 1.00 | 1.00 | 1.00 | 1.00 | 1.00 |
| Ec_PK1 | U68074 | 1.00 | 1.00 | 1.00 | 1.00 | 1.00 | 1.00 |
| T2_gene32 | X12460 | 1.00 | 1.00 | 1.00 | 1.00 | 1.00 | 1.00 |
| MMTV_gag/pro | AF033807 | 1.00 | 1.00 | 1.00 | 1.00 | 1.00 | 1.00 |
| TMGMV_UPD-PK1 | M34077 | 1.00 | 1.00 | 1.00 | 1.00 | 1.00 | 1.00 |
| STMV_UPD2-PK1 | M25782 | 1.00 | 0.92 | 1.00 | 0.92 | 1.00 | 0.92 |
| SRV1_gag/pro | M11841 | 0.92 | 0.73 | 0.92 | 0.73 | 0.92 | 0.73 |
| BSBV3_UPD-PKb | Z66493 | 1.00 | 1.00 | 1.00 | 1.00 | 1.00 | 1.00 |
| STNV1_PK2 | J02399 | 0.58 | 0.88 | 0.58 | 0.88 | 0.58 | 0.88 |
| NGF-H1 | | 1.00 | 1.00 | 1.00 | 1.00 | 1.00 | 1.00 |
| NGF-L2 | | 1.00 | 1.00 | 1.00 | 1.00 | 1.00 | 1.00 |
| NGF-L6 | | 1.00 | 1.00 | 1.00 | 1.00 | 1.00 | 1.00 |
| BMV3_UPD-PK1 | V00099 | 1.00 | 1.00 | 1.00 | 1.00 | 1.00 | 1.00 |
| TRV-PSG2_PK3 | X03686 | 1.00 | 1.00 | 1.00 | 1.00 | 1.00 | 1.00 |
| biotin-PK-AN69.1 | | 1.00 | 0.85 | 1.00 | 0.85 | 1.00 | 0.85 |
| SBRMV1_UPD-PKc | AF146278 | 1.00 | 0.92 | 1.00 | 0.92 | 1.00 | 0.92 |
| SBRMV1_UPD-PKe | AF146278 | 1.00 | 0.83 | 1.00 | 0.83 | 1.00 | 0.83 |
| MMLV_PKb | AJ299445 | 1.00 | 1.00 | 1.00 | 1.00 | 1.00 | 1.00 |
| HPeV1 | L02971 | 1.00 | 1.00 | 1.00 | 1.00 | 1.00 | 1.00 |
| PYVV3 | AJ508757 | 1.00 | 0.94 | 1.00 | 0.94 | 1.00 | 0.94 |
| IAV_PK_NS | V01104 | 0.73 | 0.40 | 0.73 | 0.40 | 0.73 | 0.40 |
| IBV_PK_NS | CY018761 | 1.00 | 0.71 | 1.00 | 0.71 | 1.00 | 0.71 |
| HAV_PK1 | M14707 | 1.00 | 0.86 | 1.00 | 0.86 | 1.00 | 0.86 |
| antiHIV1-RT_1.1 | | 1.00 | 1.00 | 1.00 | 1.00 | 1.00 | 1.00 |
| PMV_psiA | NC_002598 | 1.00 | 1.00 | 1.00 | 1.00 | 1.00 | 1.00 |
| PMV_psiB | NC_002598 | 1.00 | 1.00 | 1.00 | 1.00 | 1.00 | 1.00 |
| CMMV_psiA | NC_011108 | 1.00 | 1.00 | 1.00 | 1.00 | 1.00 | 1.00 |
| Ec_RydC | NC_000913 | 1.00 | 1.00 | 1.00 | 1.00 | 1.00 | 1.00 |
| | **Mean scores** | **0.97** | **0.93** | **0.97** | **0.93** | **0.97** | **0.93** |

Supplementary Table 1. Sensitivity and specificity scores for each sequence in the small test set (sequences containing 45 or fewer stems) using exact enumeration, REMC, and QC methods. Each method was run one time. Names derive from Pseudobase. Bolded numbers at the bottom of each column represent the mean. There are several cases where the exact solution does not have a sensitivity or a specificity of 1.0 which calls to attention limitations in the scoring function. These are cases where the lowest energy configuration found by the scoring function differs from the configuration found in nature.

| Name | EMBL Number | REMC Mean | REMC Std | QC Mean | QC Std | REMC | | QC | |
|---|---|---|---|---|---|---|---|---|---|
| | | | | | | Specificity | Sensitivity | Specificity | Sensitivity |
| GaLV | M26927 | -249.00 | 0.00 | -249.00 | 0.00 | 0.70 | 0.93 | 0.70 | 0.93 |
| BaEV | D10032 | -270.00 | 0.00 | -270.00 | 0.00 | 0.00 | 0.00 | 0.00 | 0.00 |
| STMV_UPD2-PK1 | M25782 | -117.00 | 0.00 | -117.00 | 0.00 | 0.92 | 1.00 | 0.85 | 0.92 |
| BSBV3_UPD-PKb | Z66493 | -145.00 | 0.00 | -145.00 | 0.00 | 0.73 | 0.92 | 0.73 | 0.92 |
| STNV1_PK2 | J02399 | -64.00 | 0.00 | -64.00 | 0.00 | 0.88 | 0.58 | 0.88 | 0.58 |
| NGF-H1 | | -249.00 | 0.00 | -249.00 | 0.00 | 1.00 | 1.00 | 1.00 | 1.00 |
| NGF-L2 | | -184.00 | 0.00 | -184.00 | 0.00 | 1.00 | 1.00 | 1.00 | 1.00 |
| CCMV3 | M28818 | -1608.90 | 3.21 | -1609.25 | 3.03 | 0.65 | 0.71 | 0.65 | 0.71 |
| BSMVbeta | X03854 | -511.00 | 0.00 | -511.00 | 0.00 | 0.90 | 0.84 | 0.90 | 0.84 |
| ORSV-S1_PKbulge2 | U34586 | -299.00 | 0.00 | -299.00 | 0.00 | 0.89 | 0.94 | 0.89 | 0.94 |
| PSLVbeta | M81486 | -645.00 | 0.00 | -645.00 | 0.00 | 0.89 | 1.00 | 0.89 | 1.00 |
| LRSVbeta | Z46351 | -431.00 | 0.00 | -431.00 | 0.00 | 0.86 | 0.53 | 0.86 | 0.53 |
| CoxB3 | M33854 | -144.00 | 0.00 | -144.00 | 0.00 | 1.00 | 0.52 | 1.00 | 0.52 |
| BaMV | AF018156 | -108.00 | 0.00 | -108.00 | 0.00 | 1.00 | 1.00 | 0.50 | 0.50 |
| RSV | V01197 | -1415.00 | 0.00 | -1410.50 | 7.86 | 0.76 | 0.95 | 0.76 | 0.95 |
| Rr_ODCanti | D10706 | -474.00 | 0.00 | -474.00 | 0.00 | 0.65 | 1.00 | 0.65 | 1.00 |
| SBRMV1_UPD-PKc | AF146278 | -69.00 | 0.00 | -69.00 | 0.00 | 0.92 | 1.00 | 0.92 | 1.00 |
| Hs_PrP | M13899 | -151.00 | 0.00 | -151.00 | 0.00 | 0.40 | 0.55 | 0.40 | 0.55 |
| Bt_PrP | AF117327 | -195.00 | 0.00 | -195.00 | 0.00 | 0.00 | 0.00 | 0.00 | 0.00 |
| LDV-C | L13298 | -121.00 | 0.00 | -121.00 | 0.00 | 1.00 | 0.65 | 1.00 | 0.65 |
| GLV_IRES | L13218 | -366.00 | 0.00 | -366.00 | 0.00 | 0.96 | 1.00 | 0.96 | 1.00 |
| MMLV_PKb | AJ299445 | -81.00 | 0.00 | -81.00 | 0.00 | 1.00 | 1.00 | 1.00 | 1.00 |
| ribox02 | | -980.00 | 0.00 | -980.00 | 0.00 | 0.59 | 0.69 | 0.59 | 0.69 |
| WBV | NC_008516 | -196.00 | 0.00 | -196.00 | 0.00 | 1.00 | 0.74 | 1.00 | 0.74 |
| MHV | AY700211 | -204.00 | 0.00 | -204.00 | 0.00 | 1.00 | 1.00 | 1.00 | 1.00 |
| Mm_Edr | AJ006464 | -81.00 | 0.00 | -81.00 | 0.00 | 1.00 | 0.47 | 1.00 | 0.47 |
| Hs_Ma3 | NM_013364 | -121.00 | 0.00 | -121.00 | 0.00 | 1.00 | 0.69 | 1.00 | 0.69 |
| PYVV3 | AJ508757 | -199.00 | 0.00 | -199.00 | 0.00 | 0.94 | 1.00 | 0.94 | 1.00 |
| LChV-1 | X93351 | -279.00 | 0.00 | -279.00 | 0.00 | 1.00 | 0.77 | 1.00 | 0.77 |
| CTV | U16304 | -284.00 | 0.00 | -284.00 | 0.00 | 0.95 | 0.95 | 0.95 | 0.95 |
| PMWaV-2 | AF283103 | -135.00 | 0.00 | -135.00 | 0.00 | 0.93 | 0.74 | 0.93 | 0.74 |
| IAV_PK_NS | V01104 | -140.00 | 0.00 | -140.00 | 0.00 | 0.40 | 0.73 | 0.40 | 0.73 |
| IBV_PK_NS | CY018761 | -213.00 | 0.00 | -213.00 | 0.00 | 0.71 | 1.00 | 0.71 | 1.00 |
| Bs_glmS_P3.1 | U21932 | -725.00 | 0.00 | -725.00 | 0.00 | 0.83 | 0.96 | 0.83 | 0.96 |
| VMV | NC_001452 | -299.00 | 0.00 | -299.00 | 0.00 | 0.70 | 1.00 | 0.70 | 1.00 |
| HAV_PK1 | M14707 | -108.00 | 0.00 | -108.00 | 0.00 | 0.86 | 1.00 | 0.86 | 1.00 |
| antiHIV1-RT_1.1 | | -81.00 | 0.00 | -81.00 | 0.00 | 1.00 | 1.00 | 1.00 | 1.00 |
| SAH_riboswitch | 3NPQ | -159.00 | 0.00 | -159.00 | 0.00 | 0.87 | 1.00 | 0.87 | 1.00 |
| MVEV | NC_000943 | -179.00 | 0.00 | -179.00 | 0.00 | 0.72 | 1.00 | 0.72 | 1.00 |
| SESV | AF315122 | -281.00 | 0.00 | -281.00 | 0.00 | 0.86 | 0.95 | 0.86 | 0.95 |
| PMV_psiB | NC_002598 | -117.00 | 0.00 | -117.00 | 0.00 | 1.00 | 1.00 | 1.00 | 1.00 |
| CCR5_PRF | NM_000579 | -602.00 | 0.00 | -602.00 | 0.00 | 0.32 | 0.38 | 0.32 | 0.38 |
| Ec_RydC | NC_000913 | -153.00 | 0.00 | -153.00 | 0.00 | 1.00 | 1.00 | 1.00 | 1.00 |
| TPAV_psiB | JX848617 | -100.00 | 0.00 | -100.00 | 0.00 | 1.00 | 0.67 | 1.00 | 0.67 |
| SNSV1 | NC_008708 | -567.00 | 0.00 | -567.00 | 0.00 | 0.91 | 0.97 | 0.91 | 0.97 |
| TCV1 | NC_003844 | -314.00 | 0.00 | -314.00 | 0.00 | 0.67 | 0.81 | 0.67 | 0.81 |
| BCRV1 | NC_011553 | -583.00 | 0.00 | -583.00 | 0.00 | 0.83 | 1.00 | 0.83 | 1.00 |
| PMoV3 | NC_005854 | -1016.00 | 0.00 | -1016.00 | 0.00 | 0.79 | 0.79 | 0.79 | 0.79 |
| TAMV1 | NC_003833 | -526.33 | 17.91 | -539.00 | 0.00 | 0.62 | 0.65 | 0.59 | 0.62 |
| APLPV3 | NC_003453 | -342.00 | 0.00 | -342.00 | 0.00 | 0.94 | 0.83 | 0.94 | 0.83 |
| FCiLV3 | NC_006568 | -1155.00 | 0.00 | -1155.00 | 0.00 | 0.81 | 0.95 | 0.81 | 0.95 |
| PDV3 | NC_008038 | -739.00 | 0.00 | -739.00 | 0.00 | 0.83 | 0.94 | 0.83 | 0.94 |
| | **Mean Scores** | **-361.06** | | **-361.23** | | **0.81** | **0.82** | **0.80** | **0.81** |

Supplementary Table 2. Sensitivity and specificity scores for each sequence in the large test set (sequences containing more than 45 stems) using REMC and QC methods. Each method was run ten times. Names derive from PseudoBase. Bolded numbers at the bottom of each column represent the mean.

| Name | EMBL Number | SPOT-RNA Specificity | SPOT-RNA Sensitivity | ProbKnot Specificity | ProbKnot Sensitivity | ViennaRNA Specificity | ViennaRNA Sensitivity |
|---|---|---|---|---|---|---|---|
| antiHIV1-RT_1.1 | | 0.55 | 0.40 | 1.00 | 0.73 | 0.00 | 0.00 |
| APLPV3 | NC_003453 | 0.78 | 0.54 | 0.88 | 0.86 | 0.00 | 0.00 |
| BaEV | D10032 | 0.92 | 1.00 | 0.00 | 0.00 | 1.00 | 0.50 |
| BaMV | AF018156 | 1.00 | 0.50 | 0.50 | 0.50 | 1.00 | 0.50 |
| BCRV1 | NC_011553 | 0.92 | 1.00 | 0.69 | 0.80 | 0.88 | 0.58 |
| Bs_glmS_P3.1 | U21932 | 1.00 | 1.00 | 0.74 | 0.77 | 0.79 | 0.65 |
| BSBV3_UPD-PKb | Z66493 | 0.94 | 1.00 | 1.00 | 0.75 | 1.00 | 0.71 |
| BSMVbeta | X03854 | 0.93 | 0.82 | 0.67 | 0.58 | 0.84 | 0.84 |
| Bt_PrP | AF117327 | 0.78 | 0.81 | 0.33 | 0.42 | 0.53 | 0.55 |
| CCMV3 | M28818 | 0.94 | 0.89 | 0.79 | 0.84 | 0.40 | 0.33 |
| CCR5_PRF | NM_000579 | 0.74 | 0.81 | 0.38 | 0.28 | 0.38 | 0.35 |
| CoxB3 | M33854 | 0.70 | 0.56 | 0.85 | 0.74 | 0.81 | 0.76 |
| CTV | U16304 | 0.85 | 0.74 | 0.35 | 0.40 | 0.89 | 0.74 |
| Ec_RydC | NC_000913 | 1.00 | 0.50 | 0.60 | 0.56 | 1.00 | 0.50 |
| FCiLV3 | NC_006568 | 0.88 | 0.95 | 0.91 | 0.81 | 0.66 | 0.69 |
| GaLV | M26927 | 0.67 | 0.94 | 0.00 | 0.00 | 0.48 | 0.65 |
| GLV_IRES | L13218 | 0.83 | 0.45 | 0.33 | 0.30 | 0.71 | 0.45 |
| HAV_PK1 | M14707 | 0.70 | 0.64 | 0.91 | 0.83 | 0.00 | 0.00 |
| Hs_Ma3 | NM_013364 | 0.38 | 0.50 | 0.73 | 0.69 | 0.33 | 0.42 |
| Hs_PrP | M13899 | 0.89 | 1.00 | 0.00 | 0.00 | 0.39 | 0.53 |
| IAV_PK_NS | V01104 | 0.43 | 0.39 | 0.43 | 0.55 | 0.29 | 0.30 |
| IBV_PK_NS | CY018761 | 1.00 | 1.00 | 0.40 | 0.50 | 0.75 | 0.55 |
| LChV-1 | X93351 | 0.43 | 0.31 | 0.74 | 0.77 | 0.67 | 0.62 |
| LDV-C | L13298 | 0.63 | 0.63 | 0.43 | 0.59 | 0.58 | 0.74 |
| LRSVbeta | Z46351 | 1.00 | 1.00 | 0.81 | 0.85 | 0.67 | 0.56 |
| MHV | AY700211 | 1.00 | 0.95 | 0.55 | 0.67 | 0.63 | 0.53 |
| Mm_Edr | AJ006464 | 0.89 | 1.00 | 0.59 | 0.53 | 0.73 | 0.69 |
| MMLV_PKb | AJ299445 | 0.94 | 1.00 | 1.00 | 0.55 | 0.92 | 0.69 |
| MVEV | NC_000943 | 0.77 | 0.77 | 0.36 | 0.56 | 1.00 | 0.77 |
| NGF-H1 | | 0.65 | 0.55 | 0.92 | 0.65 | 0.55 | 0.55 |
| NGF-L2 | | 0.93 | 0.74 | 1.00 | 0.88 | 0.93 | 0.74 |
| ORSV-S1_PKbulge2 | U34586 | 0.92 | 1.00 | 0.45 | 0.50 | 0.43 | 0.55 |
| PDV3 | NC_008038 | 0.67 | 0.67 | 0.62 | 0.74 | 0.46 | 0.50 |
| PMoV3 | NC_005854 | 0.82 | 0.88 | 0.73 | 0.79 | 0.91 | 0.77 |
| PMV_psiB | NC_002598 | 0.67 | 1.00 | 0.25 | 0.23 | 0.41 | 0.50 |
| PMWaV-2 | AF283103 | 0.92 | 0.92 | 0.95 | 1.00 | 1.00 | 0.58 |
| PSLVbeta | M81486 | 1.00 | 0.91 | 0.70 | 0.68 | 0.63 | 0.45 |
| PYVV3 | AJ508757 | 0.58 | 0.54 | 0.73 | 0.69 | 0.38 | 0.46 |
| ribox02 | | 0.67 | 1.00 | 0.77 | 0.85 | 0.46 | 0.61 |
| Rr_ODCanti | D10706 | 0.83 | 0.79 | 0.52 | 0.65 | 0.32 | 0.42 |
| RSV | V01197 | 0.60 | 0.46 | 0.78 | 0.82 | 0.00 | 0.00 |
| SAH_riboswitch | 3NPQ | 0.23 | 0.09 | 0.35 | 0.46 | 0.28 | 0.28 |
| SBRMV1_UPD-PKc | AF146278 | 0.94 | 1.00 | 0.80 | 0.73 | 0.82 | 0.56 |
| SESV | AF315122 | 1.00 | 0.67 | 0.32 | 0.42 | 1.00 | 0.67 |
| SNSV1 | NC_008708 | 0.71 | 0.77 | 0.71 | 0.77 | 0.83 | 0.77 |
| STMV_UPD2-PK1 | M25782 | 0.71 | 0.63 | 0.92 | 1.00 | 0.71 | 0.81 |
| STNV1_PK2 | J02399 | 0.70 | 0.77 | 0.92 | 0.92 | 0.80 | 0.80 |
| TAMV1 | NC_003833 | 0.83 | 0.85 | 0.76 | 0.86 | 0.79 | 0.79 |
| TCV1 | NC_003844 | 0.95 | 0.95 | 0.73 | 0.81 | 0.78 | 0.86 |
| TPAV_psiB | JX848617 | 0.91 | 0.86 | 1.00 | 0.87 | 0.91 | 0.86 |
| VMV | NC_001452 | 0.72 | 0.70 | 0.35 | 0.50 | 0.91 | 0.81 |
| WBV | NC_008516 | 0.78 | 0.58 | 0.54 | 0.74 | 0.74 | 0.81 |
| | | **0.79** | **0.76** | **0.63** | **0.63** | **0.64** | **0.56** |

Supplementary Table 3. Sensitivity and specificity scores for each sequence in the large test set (sequences containing more than 45 stems) using three methods found in literature. Bolded numbers at the bottom of each column represent the mean.

#### Scaling: Number of possible stems as a function of sequence length

There is no concise function that can describe the expected number of possible stems given the length of an RNA sequence. In the most limiting case, there could be 0 possible stems for a sequence of length *N*. For example, a sequence containing one nucleotide repeated *N* times, CCCC… is unable to form intramolecular base pairs. In the most extreme case, there is an exponential relationship between the number of possible stems and the length of the sequence. For example, given the following sequence: GCGCGC… there are many opportunities to form base pairs. Given a minimum stem length $L_s$, minimum loop length $L_l$, the largest possible stem would occur when the 5’ and 3’ ends for a base pair and the rest of the consecutive pairs zip up until the minimum stem length is reached:

$$L_{s,max} = \left\lfloor \frac{(N - L_l)}{2} \right\rfloor \quad (S1)$$

Similarly, the largest possible loop would occur when the 5’ and 3’ ends form a base pair and the $L_s$ neighboring bases form pairs:

$$L_{l,max} = N - 2L_s \quad (S2)$$

Therefore, the number of stems of length $L_{s,i}$ with loops of length $L_{l,j}$ is given by:

$$n_{ij} = N - \left(2L_{s,i} - L_{l,j}\right) + 1 \quad (S3)$$

By summing over all $n_{ij}$ the total number of possible stems can be computed for an RNA sequence of length *N*:

$$n = \sum_{i=L_s}^{L_{s,max}} \sum_{j=L_l}^{L_{l,max}} n_{ij} = \sum_{i=L_s}^{L_{s,max}} \sum_{j=L_l}^{L_{l,max}} N - \left(2L_{s,i} - L_{l,j}\right) + 1 \quad (S4)$$

Given an RNA sequence with $n$ possible stems, there are $2^n$ possible combinations of stems. Therefore, the total number of possible combinations of stems given a sequence of the form GCGCGC… with total length $N$ is

$$C_{GCGC\ldots} = 2^{\sum_{i=L_s}^{L_{s,max}} \sum_{j=L_l}^{L_{l,max}} N-(2L_{s,i}-L_{l,j})+1} \quad (S5)$$

In the limit where $N >> L_s$ and $N >> L_l$,

$$\lim_{\substack{N\gg L_s \\ N\gg L_l}} n \approx \sum_{i=L_s}^{\lfloor N/2 \rfloor} \sum_{j=L_l}^{N} N \approx \frac{1}{2} N^3$$

Therefore,

$$\lim_{\substack{N\gg L_s \\ N\gg L_l}} C_{GCGC\ldots} \approx 2^{N^3/2}$$

For a GCGCGC… sequence of length 1,000, there would be approximately $2^{500,000}$ possible combinations of stems. Of course, for this simple example, the global minimum is the stem referred to in Equation (S1), but the contrast between this sequence and the one where 0 stems are possible demonstrates how variable the combinatorial space can be for RNA structure prediction.

#### Direct QPU performance

D-Wave offers an API that allows users to interface directly with the QPU. The results of utilizing this API with a minimum annealing time set to 500 ms yielded the results shown in Supplementary Figure 1. Each system was run a total of 10 times on both the QPU and with the hybrid solver. The hybrid solver returned the same answer in every case. The QPU found the exact answer in all 10 trials for 20 out of 36 systems. In the other systems, particularly the larger ones, there was extreme variability in the results. Supplementary Figure 1 (right) shows 4 cases where the standard deviation exceeds the absolute value of the known solution. In each of these cases the QPU failed to return a valid solution, meaning each bitstring returned contained overlapping stems. Further studies are required to tune the system-level parameters to reduce noise and optimize the results.

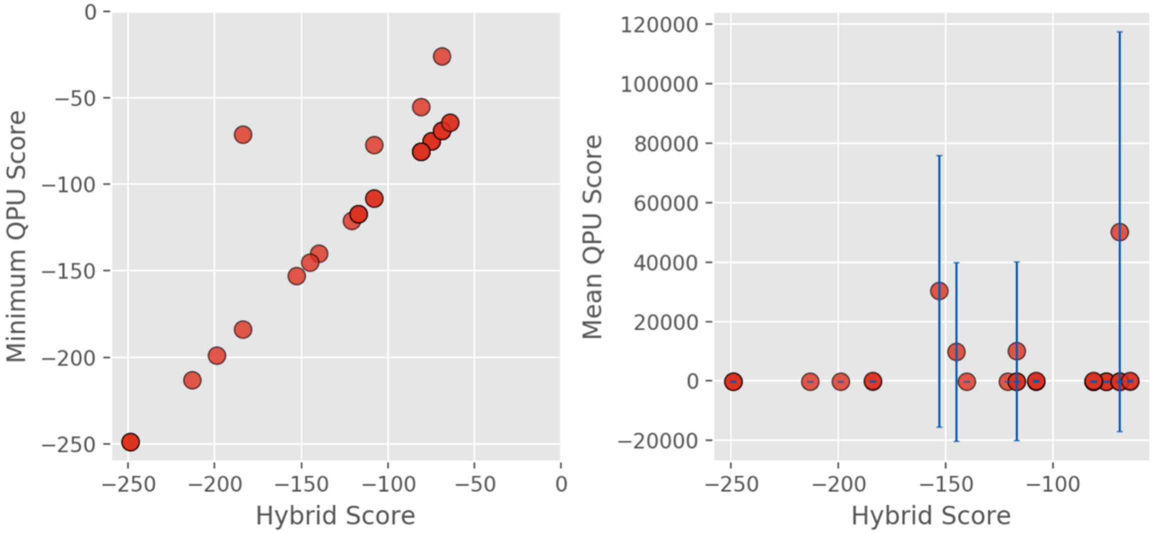


Supplementary Figure 1. (left) Minimum QPU score vs hybrid score for systems containing <60 stems. Points that fall on the line y=x indicate that the best score from the QPU is equal to the best score from the hybrid solver. (right) Average QPU score vs hybrid score. Standard deviation in the QPU scores is represented by blue bars. Positive scores indicate no suitable solution was found.

| Name | N Stems | Hybrid Score | QPU Min Score | QPU Mean Score | QPU Score Std |
|---|---|---|---|---|---|
| TMGMV_UPD-PK1 | 6 | -75 | -75 | -75.00 | 0.00 |
| BMV3_UPD-PK1 | 7 | -75 | -75 | -75.00 | 0.00 |
| TRV-PSG2_PK3 | 8 | -75 | -75 | -75.00 | 0.00 |
| CMMV_psiA | 9 | -75 | -75 | -75.00 | 0.00 |
| FIV | 14 | -81 | -81 | -81.00 | 0.00 |
| Ec_PK1 | 14 | -81 | -81 | -81.00 | 0.00 |
| HPeV1 | 14 | -81 | -81 | -81.00 | 0.00 |
| PMV_psiA | 14 | -81 | -81 | -81.00 | 0.00 |
| BSBV3_UPD-PKb | 16 | -108 | -108 | -108.00 | 0.00 |
| MMLV_PKb | 16 | -81 | -81 | -81.00 | 0.00 |
| T2_gene32 | 17 | -69 | -69 | -69.00 | 0.00 |
| NGF-H1 | 17 | -249 | -249 | -249.00 | 0.00 |
| NGF-L6 | 18 | -249 | -249 | -249.00 | 0.00 |
| biotin-PK-AN69.1 | 18 | -117 | -117 | -117.00 | 0.00 |
| SBRMV1_UPD-PKe | 21 | -69 | -69 | -69.00 | 0.00 |
| STNV1_PK2 | 23 | -64 | -64 | -64.00 | 0.00 |
| SRV1_gag/pro | 24 | -121 | -121 | -121.00 | 0.00 |
| NGF-L2 | 24 | -184 | -184 | -176.20 | 23.40 |
| HAV_PK1 | 24 | -108 | -108 | -108.00 | 0.00 |
| IBV_PK_NS | 27 | -213 | -213 | -209.80 | 9.60 |
| PMV_psiB | 27 | -117 | -117 | -111.20 | 11.60 |
| SBRMV1_UPD-PKc | 28 | -69 | -69 | -69.00 | 0.00 |
| IAV_PK_NS | 28 | -140 | -140 | -138.13 | 4.96 |
| antiHIV1-RT_1.1 | 29 | -81 | -81 | -75.75 | 13.89 |
| MMTV_gag/pro | 30 | -69 | -69 | -60.13 | 15.82 |
| PYVV3 | 30 | -199 | -199 | -179.75 | 19.26 |
| STMV_UPD2-PK1 | 32 | -117 | -117 | -104.88 | 23.15 |
| Ec_RydC | 45 | -153 | -153 | 30288.10 | 45748.96 |
| NGF-H1 | 45 | -249 | -249 | -147.13 | 55.95 |
| STNV1_PK2 | 46 | -64 | -64 | 151.70 | 185.67 |
| BSBV3_UPD-PKb | 48 | -145 | -145 | 9965.10 | 30005.69 |
| MMLV_PKb | 48 | -81 | -55 | -7.40 | 32.02 |
| HAV_PK1 | 51 | -108 | -77 | 210.50 | 219.73 |
| NGF-L2 | 52 | -184 | -71 | 7.40 | 58.04 |
| SBRMV1_UPD-PKc | 59 | -69 | -26 | 50206.60 | 67155.91 |
| STMV_UPD2-PK1 | 59 | -117 | -117 | 10151.70 | 29958.26 |

Supplementary Table 4. Direct programming QPU vs Hybrid solver.
